# Longevity and environmental temperature modulate mitochondrial DNA evolution in fishes

**DOI:** 10.1101/2024.03.07.583929

**Authors:** Sabrina Le Cam, Mathieu Mortz, Pierre Blier

**Author notes:** First coauthors (Sabrina Le Cam, Mathieu Mortz).

## Abstract

The link between longevity and mitochondrial function has been documented for years. Since mitochondrial DNA (mtDNA) encodes for electron transport system (ETS) proteins, we could suspect that its evolution is linked with that of longevity. A negative relationship has been documented between the synonymous substitution rate and lifespan when analyzing the whole mitochondrial genome in animals. In this study, we aimed to confirm this negative correlation for each of the mitochondrial protein coding genes (mtPCGs) and explore potential relationships between adaptation to extreme temperatures and the evolution of mtDNA. To this end, we selected 112 species of fish with a wide range of longevity as well as divergences in environmental temperature, which is a good proxy for energy metabolism in these animals. Our results 1) challenge the “rate of living” theory by not showing any correlation between longevity and environmental temperature, 2) confirm the negative relationship between substitution rate and longevity for each of the 13 mtPCGs, and 3) highlight for the first time a link between high conservation of the three COX genes and adaptation to warmer temperatures in fish. By challenging a paradigm and extending the conclusions made for mtDNA to individual genes, our study opens a wide field to be explored concerning study of the aging process. Moreover, the specific link between the evolution of COX genes and temperature tolerance confirms the importance of complex IV in adaptation to extreme temperatures and, more generally, the importance of distinguishing gene families when studying mtDNA evolution in animals.

## Introduction

Understanding the mechanisms behind aging has become a major issue of the 21^st^ century. Among the hallmarks of aging processes (*Bratic et al. 2013; Sun et al. 2016; Son et al. 2019*), mitochondrial dysfunction is thought to be a key factor, but it remains to be determined if this impairment is at the origin or a consequence of aging. The “free-radical theory of aging” formulated by Harman in the 1950s (*Harman 1956*) advocated that free radicals and reactive oxygen species, naturally produced in cells supported by aerobic metabolism, could harm surrounding biomolecules. It was thought that the accumulation of defective proteins, DNA, or lipids would ultimately lead to the collapse of homeostasis.

This naive perception of the management of oxidative stress and its impact on lifespan has been fiercely criticized, which led to a theory more focused on mitochondria (mitochondrial oxidative stress theory of aging, MOSTA) (*Scialo et al. 2013; Blier et al. 2017*). MOSTA speculates that mitochondria are not only a major source of cellular ROS but also the principal target of these ROS. Indeed, ROS principally produced by the electron transport system (ETS) can react with essential biomolecules, particularly mitochondrial inner-membrane lipids (*reviewed in Hulbert et al. 2008; Munro and Blier 2012*), leading to several cellular dysfunctions. The cumulative effect of this biochemical damage causes mitochondrial DNA oxidation or mutations. Alterations in mtDNA are commonly observed during aging processes, causing functional impairments on mitochondrial protein coding genes (mtPCGs) (*Agarwal et al. 1995*) and contributing to the decline of the mitochondrial respiratory function (*Wei et al. 2009*). Thus, the accumulation of mutations on mtDNA has been already linked to degenerative diseases and longevity in vertebrates (*Trifunovic et al. 2004*). Therefore, if mitochondrial functions and their decline exert any control on the aging process and species lifespan, one can hypothesize that the pattern or mode of mitochondrial DNA evolution should reveal marks of selection associated with lifespan.

It is on this basis that some researchers explored the existing relationship between the mtDNA substitution rate and the individual lifespans in fishes, mammals, and birds (*Galtier et al. 2009a, b; Hua et al. 2015*). These studies revealed a negative relationship between the mtDNA substitution rate and longevity, suggesting the need to maintain the structural integrity of mitochondrial-encoded proteins, particularly in long-lived species, to maintain their biological function. In a recent publication (*Mortz et al. 2021*), we confirmed this observation for a representative invertebrate group when we observed a negative correlation between the mtDNA synonymous substitution rate (dS) and longevity in a bivalve species. To complement all these studies, we wanted to take a more precise approach by exploring each mitochondrial gene in a phylogenetic survey of mtDNA evolution. Until now, mtDNA has been studied in its entirety, and no study has examined the relationship between phenotypic traits like longevity and the substitution rate of individual mitochondrial genes. We therefore wanted to separately investigate each of the 13 mitochondrial mtPCGs, which we grouped by family (COX and ND) to increase the statistical power of the analysis. Knowing that a decrease in complex I activity has been associated with increased lifespan (*Miwa et al. 2014*), we predicted that the rate of evolution of genes from this complex (ND family) would be particularly linked with the maximum lifespan recorded from different species.

We decided to focus on fish species, partly because of their great diversity (more than 20 000 species) and their wide range of lifespan (*Das 1994; Patnaik et al. 1994*). Many aging studies use fish as a biological model, like the zebrafish (*Danio rerio*) (*Kishi 2004; Van Houcke et al. 2015; Carneiro et al. 2016; Barth et al. 2019*) and other short-lived species (*Žák et al. 2021*) such as fishes of the genus *Nothobranchius* (*Genade et al. 2005; Platzer and Englert 2016; Hu and Brunet 2018; Barth et al. 2019*). Another major interest in studying ectotherms (fish) instead of endotherms (birds and mammals) is the possibility to test the “rate of living theory” of aging, which postulates that lifespan is dictated by the organism’s metabolic rate. Considering that environmental temperature is associated with metabolic rate, this hypothesis predicts that lifespan should be inversely correlated to the environmental temperature tolerated by fish, which would also be related to the pattern of mtDNA evolution. As part of the study reported here, we recovered from the literature the longevity records of 112 fishes that we supplemented, when possible, with the generation time, morphological traits (length and weight), and environmental traits (minimal and maximal temperature of their natural habitat) (Table S1). The aim was therefore to determine whether trait changes were associated with changes in mtPCGs during evolution, and in this way to highlight specific mtDNA modifications correlated with increased or decreased longevity, or with other life-history traits. For this purpose, we applied a Bayesian phylogenetic reconstruction and Markov chain Monte Carlo method developed by Lartillot and Pujol (*2011*), which has already been successfully used in numerous studies (*Galtier et al. 2009a, b; Nabholz et al. 2011, 2013; Mortz et al. 2021*). With this method, we were able to estimate synonymous substitution rates (dS) and the ratio between the non-synonymous and synonymous substitution rates (dN/dS), and their correlations with phenotypic traits (*Lartillot and Pujol 2011*).

Here we describe for the first time the existing relationships between the evolution of each mitochondrial gene and six phenotypic traits (generation time, longevity, length, weight, minimal and maximal temperature) in fish species. The differences highlighted here between COX and ND genes confirms that a distinction must be made between mtPCGs when studying mtDNA evolution in the future.

## Results / Discussion

### 1) Phylogeny of fishes and relationships among life-history traits

As a starting point, we needed to establish the phylogeny between the 112 selected fish species. From the concatenation of all mt-PCGs and of 12S and 16S rRNA gene alignments, we made two distinct maximum-likelihood phylogenetic trees (Figure 1), then we chose the tree based on rRNAs alignments which represents in a more discriminating way the genetic relationships among bony fishes that are already described in the literature and based on nuclear and mitochondrial DNA sequences *(Betancur et al. 2013; Satoh et al. 2016; Betancur et al. 2017*) or expressed sequence tags (*Oleksiak 2010*). Overall, we obtained nine groups of fishes (Figure 1), with the ancestral order of Petromyzontiformes, the cartilaginous fishes belonging to the class of Chondrichthyes, and seven orders of bony fishes (Osteichthyes).

**Figure 1.**
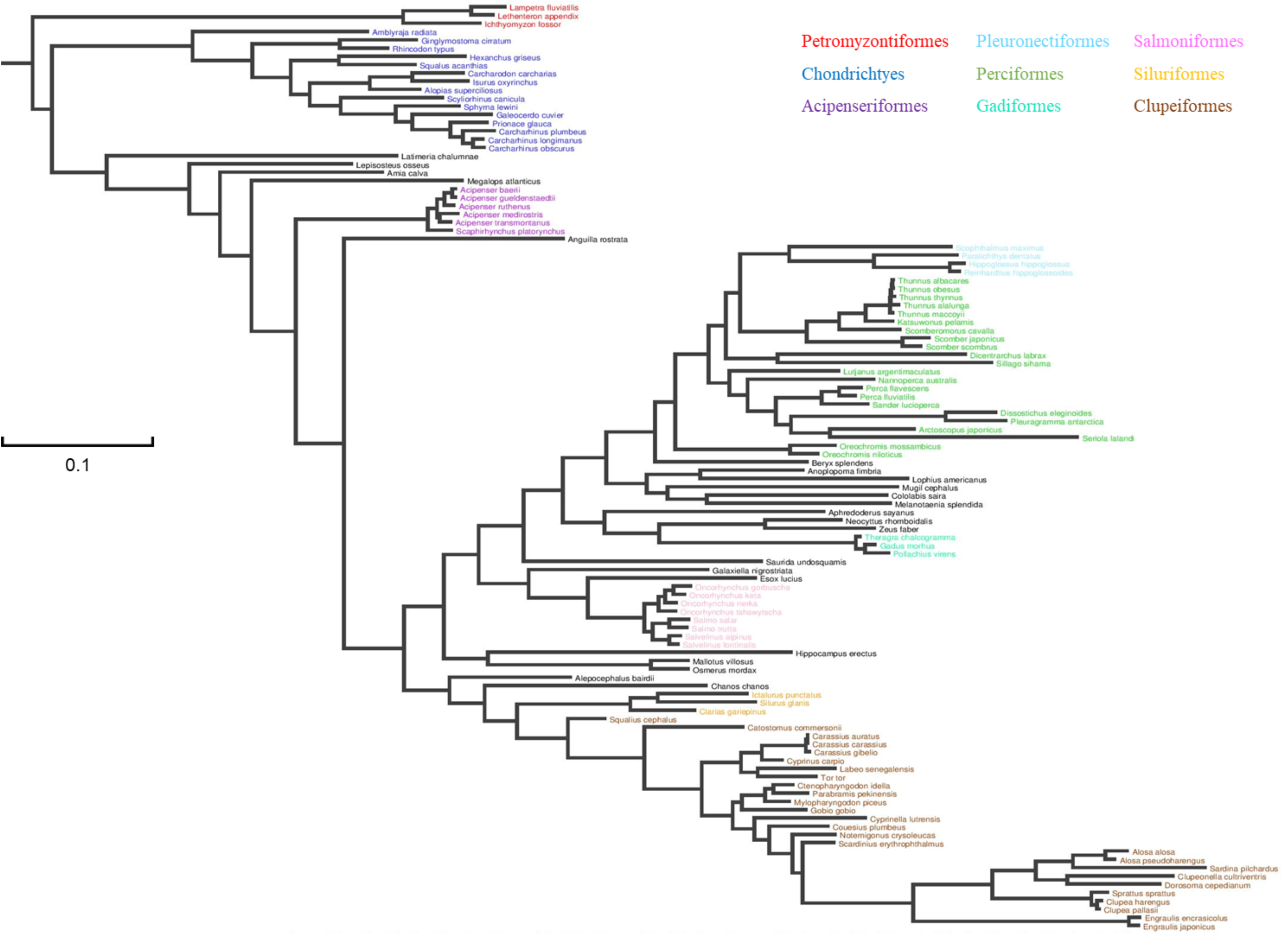
– Phylogenetic tree of 112 fish es obtained by the maximum-hood method using the site-ogenous CAT-GTR model.

As the selection of species is very dependent on the available sequences and data for these fish, we are aware that this phylogenetic representation may not be the most adequate. However, as the coevolution analysis presented below considers the phylogenetic distance between species, the choice of groups does not bias the results presented, even if they cannot be validated for the groups of fishes absent from the analysis. The Acipenseriformes, a primitive order of Actinopterygii, are more easily distinguishable, notably followed by six orders of class Euteleostei: the Clupeiformes and Siluriformes, belonging to the subclass of Otocephala, the Salmoniformes (Protacanthopterygii), the Gadiformes (Paracanthopterygii), and the Acanthopterygii orders Perciformes and Pleuronectiformes. This phylogeny was then used for correlation analysis between life-history traits, and between life-history traits and mtDNA gene substitution rates. By analyzing the covariances between the studied phenotypic traits, we were able to observe the correlation between phenotypic traits, which have generally been known and established for years, for the 112 fish species that we selected for this study. In addition, the acknowledged interdependencies between the different life-history traits selected here allowed us to corroborate the validity of the measures collected for the analysis. Thus, the allometric relationships between longevity, generation time, and species length and weight are easily found, as shown by the linear relationships between the logarithm of these measures (Figure 2 A–F). As expected, there is a positive relationship between longevity and generation time (R^2^=0.40), organism length (R^2^=0.52), and organism weight (R^2^=0.33), in agreement with the “rate of living theory,” which links an organism’s metabolic rate to its lifespan (*Liochev 2013; Scialo et al. 2013*). Larger animals have a lower energy metabolism per unit mass and, according to the “rate of living theory,” should have a higher life expectancy than smaller species (*Speakman 2005*).

**Figure 2.**
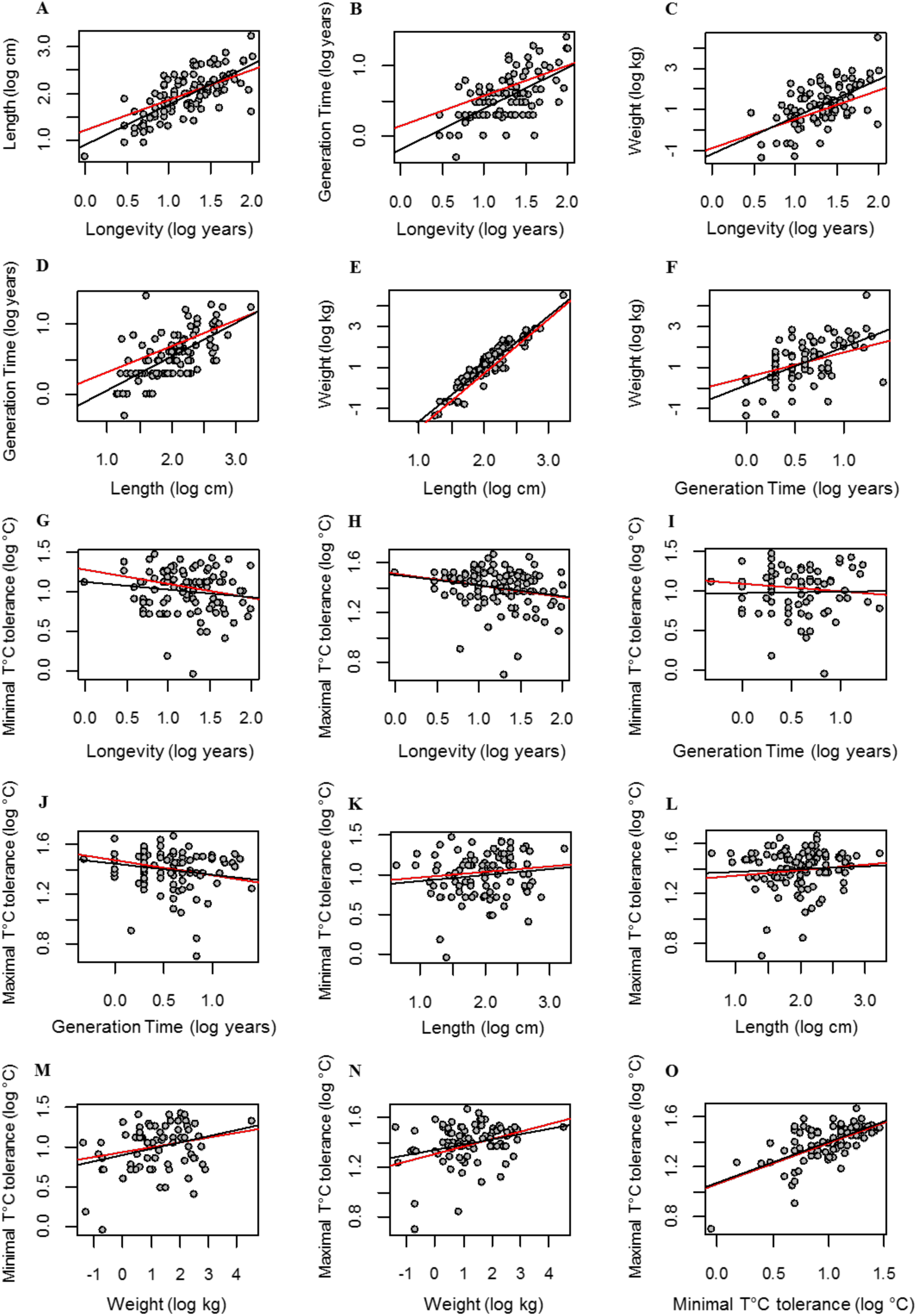
– Covariance analysis between life-history traits in 112 fish species, by following a linear model (black slope) and considered the covariance structure due to phylogenetic relations between species (red slope).

In ectothermic organisms such as fishes, habitat temperature should have an important impact on life-history traits such as the metabolic rate (*Gilloly et al. 2001*) and longevity (*Klass 1977*; *Lee and Kenyon 2009; Mori and Sasakura 2009*), with higher metabolic rates and a shorter life expectancies for animals at high temperatures. The BOLT (Burn Out Lifespan and Thermosensivity) theory suggested by Bolt and Bergman in 2015 presents a good review of the harmful effects of high temperatures on cell physiology, and even goes so far as to assert that heat and thermosensitivity are the main factors responsible for aging (*Bolt and Bergman 2015*). Higher temperatures will induce modifications in the topologies of biomolecular networks as well as of their properties, leading to acceleration of the cellular aging processes (*Bolt and Bergman 2015*). It has also been shown that animals exposed to colder temperatures have reduced standard metabolic rates and costs of living, allowing them to allocate more energy for growth; this suggests a possible negative correlation between body size and habitat temperature (*Killen 2014*). However, we could establish no relationship between the critical thermal maximum (CTmax) or minimum (CTmin) of studied fishes and the other phenotypic traits (Figure 2 G– N), which are longevity (Figure 2 G–H), and length (Figure 2 K–L). It is not possible to identify mechanisms that would compensate for the deleterious effects of high temperatures on organisms, and the literature on this subject does not provide unambiguous answers.

When comparing fish populations, Uliano et al. (*2010*) noted that those living in tropical areas have a better ability to limit the increase in energy metabolism in response to an increase in temperature than those living in temperate and subtropical environments. However, it is the increase in energy metabolism at higher temperatures (*Clarke et al. 1999*) that seems to be responsible for the decrease in life expectancy in animals. The thermal adaptations of species living in extreme temperature environments could therefore explain the lack of correlation between longevity and temperature for studied fishes. In any case, the deleterious effects of oxidative stress—caused by an overproduction of ROS, itself dependent on energy metabolism and temperature—do not seem to be the only ones responsible for premature aging in fish.

Mitochondrial structure and functions could also be modulated through membrane composition to accommodate environmental temperature and longevity (the membrane-pacemaker theory of aging; *Hulbert et al. 2008*). In agreement with MOSTA (*Scialo et al. 2013*; *Blier et al. 2017*), this theory emphasizes the importance of membrane composition to protect against the risk of oxidation by ROS and reactive carbonyl species, which are at the origin of cellular dysfunctions leading to aging processes. The likelihood of membranes being oxidized will greatly depend on their content of polyunsaturated fatty acids (PUFA), which are less resistant to peroxidation than saturated (SFA) or monounsaturated (MFA) fatty acids (*Holman, 1954*), and can be estimated by determining the peroxidation index (PI). Thus, a negative correlation between PI and lifespan has been demonstrated in clades of vertebrate (*Hulbert et al. 2007; Valencak and Ruf 2013; Pamplona et al. 2008*) and invertebrate (*Munro and Blier 2012*) animals, with a decrease in the ratio of PUFA associated with greater longevity. Membrane fatty acids composition will notably depend on body size (*Couture and Hulbert 1995*), with a decrease in PUFA levels in larger animals that may explain their longer lifespan. The PI will also be strongly dependent on temperature, especially in ectothermic organisms such as fish, where we observe important remodeling of the membrane lipid composition during thermal acclimation (*Snyder et al. 2012; Fadhlaoui et al. 2018; Malekar et al. 2018*). This process is called homeoviscous adaptation (*Hazel 1995*), and it seems to mainly affect mitochondrial membranes (*Hazel and Williams 1990*). Exposure of fish to colder temperatures would lead to increased PUFA levels and could therefore cause a decrease in life expectancy (*Guderley 2004; Snyder et al. 2012*); in contrast, acclimatization or adaptation to higher temperatures decreases PUFA levels and allows better resistance of the biological membranes to the aging processes (*Malekar et al. 2018*). We can therefore assume that these processes could counteract each other to explain the absence of a link between temperature and longevity in our analysis. However, if mitochondrial membrane composition counteracts the impact of higher temperatures on metabolism and lifespan, then metabolic rate would not be the main driver of longevity, thereby refuting the “rate of living theory,” at least in fish.

### 2) Correlation between mtPCG evolution and life-history traits: differences between COX and ND genes?

In addition to being the target of ROS during oxidative stress, mitochondria are also a main production site of these reactive species, highlighting the expected involvement of mitochondrial ETS complex activity in ROS-induced senescence processes. The sites at which most of the ROS production is observed are complexes I and III, for which some protein subunits are encoded by the mitochondrial genome. It is currently known that the evolution of mitochondrial DNA is linked to certain life-history traits such as longevity, with a negative correlation between the rate of synonymous substitution of mtPCGs and life expectancy in fishes, mammals, and birds, (*Galtier et al. 2009a, b; Hua et al. 2015*) and for invertebrate bivalves (*Mortz et al. 2021).* Here we further analyze the relationship between substitution rate and longevity by distinguishing between the two groups of genes coding for subunits of complex I (ND genes) and IV (COX genes) (Tables 1 and 2), and also by replicating the analysis individually for each of the 13 mitochondrial genes (Figure 3). The global analysis done on each of the two blocks of genes (COX and ND) reveals interesting disparities to explore.

**Figure 3.**
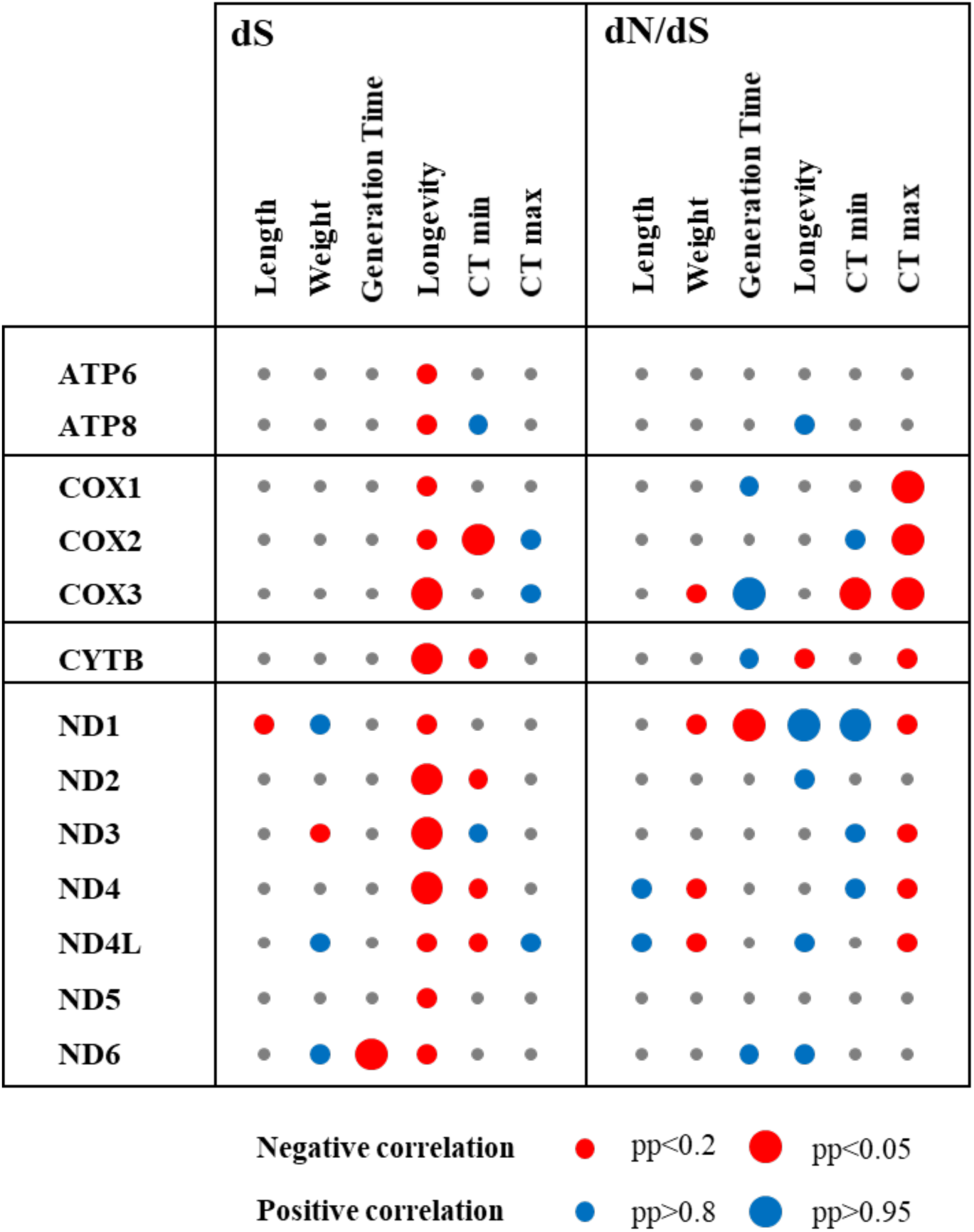
– Covariance between the synonymous substitution rate (dS) and the ratio of the non-synonymous and the synonymous substitution rates (dN/dS) of all individual mitochondrial genes and life-history traits for 112 fish species. Circle colours indicate whether the correlation between gene evolution and the phenotypic trait was positive (blue), negative (red), or absent (gray). Circle size indicates the degree of significance of the relationship highlighted, with small circles representing pp<0.2 or >0.8 and large circles representing pp<0.05 or >0.95.

**Table 1.** – Covariance between the synonymous substitution rate (dS) of mitochondrial concatenate COX and concatenate ND genes (three and seven genes respectively) and life-history traits for 112 fish species. Significant correlations are shown in bold (posterior probability (pp) <0.05; pp>0.95).

|  | COX genes |  |  | ND genes |  |  |
| --- | --- | --- | --- | --- | --- | --- |
|  | r | r<sup>2</sup> | pp | r | r<sup>2</sup> | pp |
| Length | -0.062 | 0.004 | 0.360 | -0.185 | 0.034 | 0.110 |
| Weight | 0.088 | 0.008 | 0.700 | 0.081 | 0.007 | 0.710 |
| Generation Time | -0.022 | 0.000 | 0.460 | -0.039 | 0.002 | 0.400 |
| Longevity | **-0.370** | **0.137** | **0.022** | -0.200 | 0.040 | 0.080 |
| CT min | -0.115 | 0.013 | 0.270 | **-0.283** | **0.080** | **0.034** |
| CT max | 0.065 | 0.004 | 0.640 | 0.038 | 0.001 | 0.610 |

**Table 2.**
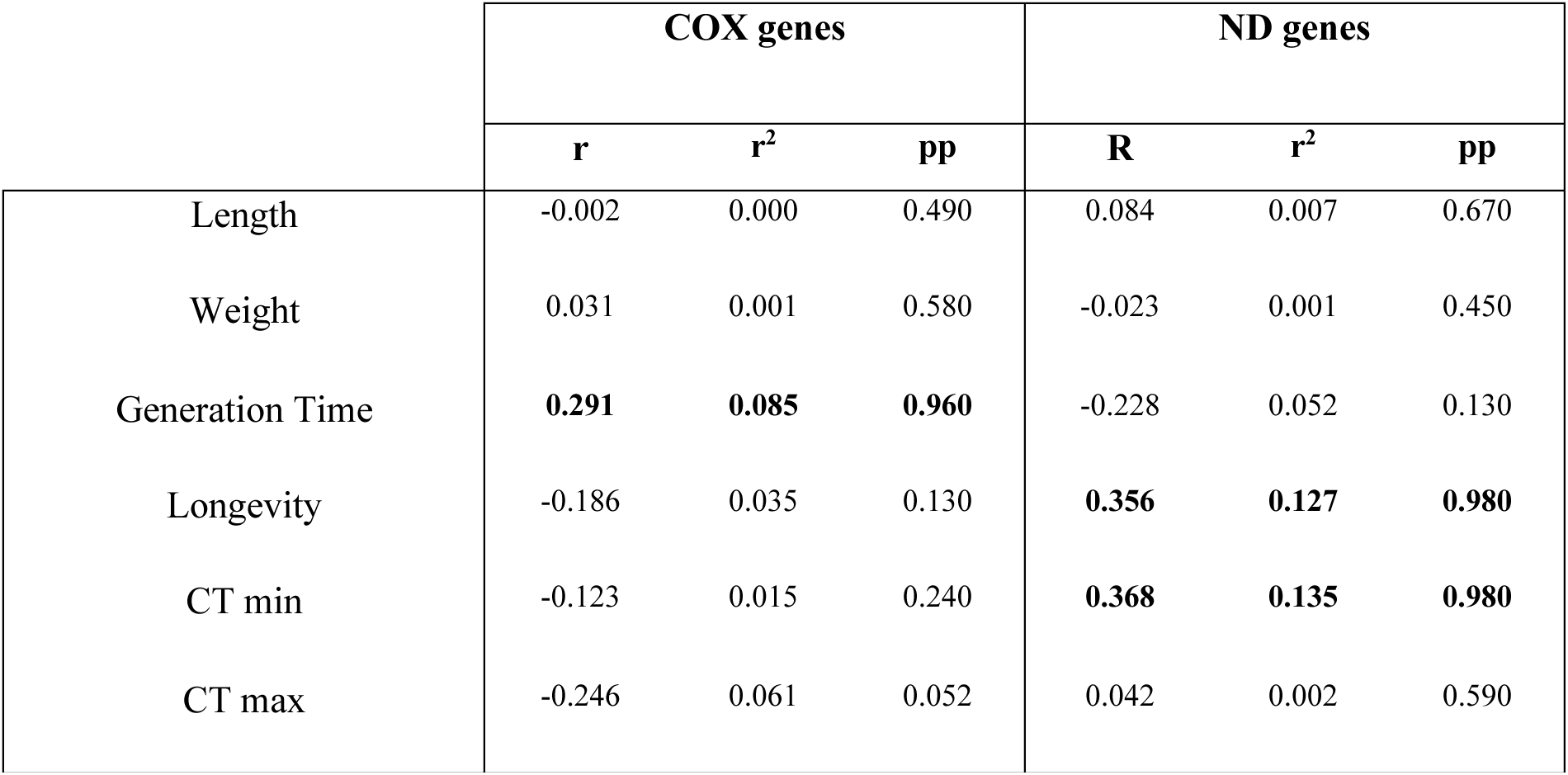
– Covariance between the ratio of the non-synonymous and the synonymous substitution rates (dN/dS) of mitochondrial concatenate COX and concatenate ND genes and life-history traits for 112 fish species. Significant correlations are shown in bold (pp<0.05; pp>0.95).

We confirmed the existence of a negative correlation between the synonymous substitution rate and longevity for each of the two groups of genes studied; the correlation was much stronger for COX genes (r=–0.37; pp=0.02) than for ND genes (r=–0.20; pp=0.08) (Table 1). However, many observations have emphasized the importance of complex I (composed of subunits encoded by ND genes) in the regulation of aging through two independent mechanisms involving either the production of ROS or the control of the NAD+/NADH ratio (reviewed by *Stefanatos and Sanz 2011* and *Scialo et al. 2013*). In addition, it has been shown that a defective assembling of complex I could have a negative effect on longevity in mice (*Miwa et al. 2014*) in the same way that insects are affected with a complex I inhibitor treatment such as tebufenpyrad (*Jovanovic et al. 2014*). Concerning respiratory chain complex IV—also called cytochrome c oxidase (CytOx) and composed of 13 subunits, among which three are encoded by the mitochondrial genes COX I, II, and III—a link between the maintenance of its activity and longevity has been demonstrated in Drosophila (*Klichko et al. 2014*). In addition, it should be noted that mutations in the COX III gene will also lead to an increase in ROS production and a decrease in lifespan in mice (*Reichart et al. 2019*).

In parallel, our analysis also revealed a surprising positive correlation between the dN/dS ratio of ND genes and longevity (r=0.36; pp=0.98) (Table 2), suggesting that positive selection pressure would be associated with increased longevity in fish. This relationship was not observed with COX genes (r –0.19; pp=0.13) (Table 2) and would therefore require further study to identify variants of complex I subunits encoded by the mitochondrial genome and associated with longevity in fishes. The second main result concerns the correlations between CTmin and the synonymous substitution rate (negative correlation; r=–0.28; pp=0.034) (Table 1) or the dN/dS ratio (positive correlation; r=0.37; pp=0.98) (Table 2) of ND genes, exactly like the relationships observed with longevity in these fish. This suggests that a purifying selection pressure would be associated with CTmin for ND genes, with a higher synonymous substitution rate and a lower dN/dS at lower temperatures, and therefore relaxation of selective pressure at lower temperatures. Until now, functional links between complex I activity and water temperature had been demonstrated in fish, suggesting that an increase in temperature would induce a decrease in the activity of complex I and vice versa (*Loskovich et al. 2005; Sappal et al. 2015; Onukwufor et al. 2016; Eya et al. 2017*), but none of these studies examined the influence of temperature on the evolution of mitochondrial genes coding for complex I subunits. To our knowledge, only one study has linked temperature sensitivity of mitochondrial respiration with mitochondrial haplotype divergences, and that was in *Drosopohila simulans* (*Pichaud et al. 2012*). In the same way as for the relationships highlighted with longevity, this would also merit further investigation with the aim of linking and understanding the functional consequences of the sequence differences observed. Among the other statistically significant results, some are difficult to interpret, such as the positive correlation between the dN/dS ratio of COX genes and generation time. To further explore these relationships and try to understand the observed patterns, we hypothesized that there might be differences between individual genes within the gene blocks studied.

### 3) Correlation between life-history traits and mtPCG evolution at the individual gene level

Results of the individual gene analysis are shown in Figure 3. At first glance, one can immediately notice that the nature of the complex to which the coded subunit belongs will not be the only determinant of the evolutionary process that the gene will undergo, and that correlations will then be observed with certain phenotypic traits. The negative correlation, between the mtDNA synonymous mutation rate and organism longevity, is the only one found for all the genes studied, with posterior probabilities (pp) of 0.2 and lower. In view of the pp reported by the analysis, this negative correlation seems stronger for the COX3, Cytb, ND2, ND3, and ND4 genes (pp<0.05, Figure 3). For the first time, we thus confirm for each of the 13 mitochondrial genes a result already known for the whole mitochondrial genome. For the genes ATP6, ATP8, and Cytb, no other strong correlation was observed in our analysis (Figure 3).

The evolution of these genes does not seem to be reflected in changes in the phenotypic traits studied. Among the significant relationships highlighted by our study, the one between the complex IV (or CytOx) and the CTmin and CTmax is particularly interesting. A strong negative correlation (pp<0.05) between the dN/dS of COX genes and CTmax was initially observed (Figure 3), as had been detected in the gene block analysis (Table 2). This robustly confirms that the purifying selection pressure of COX genes is associated with an increase in CTmax in fish. For the relationships highlighted between dN/dS and CTmin, there was a notable difference between the COX2 and COX3 genes. Indeed, CTmin is negatively correlated with dS but positively correlated with dN/dS of the COX2 gene, implying that a purifying selection pressure of this gene would accompany a decrease of the CTmin in fish. Conversely, the strong negative correlation between CTmin and the dN/dS of COX3 is rather indicative of relaxed purifying selection pressure of the COX3 gene in response to a decrease in CTmin. Thus, adaptation to higher temperatures seems to have the same effect on the evolution of the three COX genes contrary to adaptation to lower temperatures, which reveals differences between COX2 and COX3. A functional explanation is difficult to provide here, but it would seem that the strength of control of mitochondrial respiration by CytOx is affected by temperature changes and entails adjustment during thermal adaptation. In an analysis of the thermal sensitivity of mitochondrial respiration and of different enzymes and complexes in various tissues of six fish species and one endotherm, Blier et al. (2014) found a strong correlation between mitochondrial respiration (state 3) Q10s with CytOx Q10s, leading them to suspect strong control of respiration by CytOx over a wide range of temperatures. An increase in CytOx activity during cold acclimatation has been noted in killifish (*Grim et al. 2010; Dhillon and Schulte 2011*), in drosophila (*Pichaud et al. 2010, 2011*), or after exposure to extreme temperatures in daphnids (*Kake-Guena et al. 2017*). In drosophila, the CytOx seems to exert higher control of the mitochondrial metabolism at high temperature (*Pichaud et al. 2010, 2011*). In response to an increase in body temperature in an endothermic organism such as the rat, an increase in metabolic activity has also been shown to be due in part to an increase in CytOx content (*Mitchell et al. 2002*). Another study that looked at the activity of each complex of the respiratory chain showed a predominance of complex IV activity during cold adaptation, and conversely, of complexes I, II, and III in warm environments (*Hunter-Manseau et al. 2019*). These results confirm that the architectures of mitochondria and furthermore of ETS are dependent on the environmental temperature to which fish are adapted. Since thermal sensitivities of the various enzymes and complexes are quite variable, any change in the environmental temperature will necessarily perturb this architecture and require new adjustments.

Finally, there are other significant results emerging from this analysis that are more difficult to explain and would require further exploration. First, there is a strong correlation between generation time and the synonymous substitution rate of the ND6 gene and the dN/dS of the COX3 gene (Figure 3), implying that there is positive selection pressure for these two genes accompanying the increase in generation time. We would have instead expected relaxed selection for short generation times in most cases, which makes this result surprising and may open future perspectives concerning the link between ND6 and COX3 gene function and reproductive success in these fish. The other interesting aspect of our results concerns the ND1 gene, for which we note many strong correlations that are not found in the other genes coding for complex 1 subunits. First, we note that the positive correlation between dN/dS and longevity found for the ND gene block (Table 2) seems to be mainly due to the ND1 gene, implying that positive selection pressure would be associated with increased longevity in fish. Paradoxically, a negative correlation between generation time and the dN/dS ratio of the ND1 gene was observed in parallel, revealing negative selection pressure associated with an increase in generation time. Thus, we would have a negative selection pressure associated with a long generation time and then a relaxed purifying selection with an increase in longevity, revealing the particular importance of the ND1 gene compared to other genes in the study of the links between the evolution of mitochondrial DNA and longevity in animals. It would therefore be interesting to further investigate this subject to search for variants specifically associated with species possessing atypical phenotypic traits, such as very high longevity.

## Conclusion

In this study, we present the first study investigating the relationships between phenotypic traits and mitochondrial DNA evolution by distinguishing the 13 coding genes. Our findings are consistent with what has been previously demonstrated in whole mtDNA studies (*Galtier et al. 2009a, b; Hua et al. 2015; Mortz, Levivier et al. 2021*): there is a negative correlation between synonymous substitution rate and longevity for each of the mtPCGs in fish. By introducing tolerance temperatures into our analysis, we were also able to discuss the lack of a relationship between CTmin and CTmax temperatures and longevity. In ectothermic organisms such as fish, whose energetic metabolism is very dependent on water temperature, the “rate-of-living” theory led us to predict a decrease in longevity accompanying acclimation to warm temperatures, and vice versa. There are other mechanisms that come into play to allow fish to colonize environments outside their optimal temperature range without impacting their longevity. Finally, our analysis also detected for the first time a strong negative correlation between the dN/dS ratio of the three COX genes and CTmax in fish, highlighting the importance of complex IV in the acclimation of fish to warm temperatures. All these results open interesting perspectives linking the evolution of different mtDNA genes with phenotypic traits, revealing once again that the proper functioning of vital biological processes relies on these proteins, which therefore need to be strongly conserved during acclimatization to an extreme environment to avoid activating senescence mechanisms responsible in the longer term for organism aging.

## Methods

The complete mitochondrial genomes of 112 fish species were downloaded from GenBank and used for our analyses (Table S1). Six life-history traits were tested: length, weight, generation time, longevity, and the critical thermal minimum (CT min) and maximum (CT max). We found length, longevity, and CT max data for all 112 fish species, weight data for 89 species, generation time for 103 species, and CT min for 95 species. The references used to collect these data are indicated in Table S3. Each 13 mtPCG and the two ribosomal DNA from the 112 fish mtDNAs were aligned separately using MUSCLEv3.8.31 *(Edgar 2004),* and the poorly aligned positions were removed using Gblocks v.0.91b *(Castresana 2000)* with default parameters. A phylogenetic tree was then inferred using PhyML (*Guindon et al. 2010*) on SeaView v4.6.2 (*Gouy et al. 2010*) obtained by the maximum-likelihood method through the site-heterogenous CAT-GTR model, to increase robustness against long-branch attraction artifacts in the presence of mutational saturation (*Lartillot et al. 2007; Lartillot et al. 2013*). To obtain the most representative tree of the phylogenetic relationships among fish species already established in the literature, we tested three alignments (mtPCG concatenate, COX1, and 12S/16S concatenate) and chose to keep the one made from a concatenate of the genes encoding 12S and 16S rRNA. The Petromyzontiformes species were used as an outgroup for phylogenetic analyses. The correlation between life-history traits were analyzed using R 4.1.0, and the covariance structure due to phylogenetic relations between species was estimated through the R package phytools (*Revell 2012*)., The correlation and covariation among life-history traits and mtDNA evolution were then measured using a Bayesian framework with a Markov chain Monte Carlo (MCMC) sampling approach as implemented in CoEvol 1.4b (*Lartillot and Pujol 2011*). The strength of the correlation between the evolution rate and life-history traits is given as a posterior probability (*pp*) of a positive (*pp* close to 1) or a negative (*pp* close to 0) correlation. The mtDNA evolution was represented here by considering the synonymous substitution rate (dS) and the ratio of nonsynonymous to synonymous substitution rates (dN/dS). Two independent runs were performed for each analysis during 3200 cycles, excluding the first 300 cycles (burn-in). Their convergence was assessed by measuring several key statistics (e.g., log likelihood, mean substitution rate over the tree, mean omega over the tree, entries of the covariance matrix, root age), the effective sample size, and the discrepancy between the credibility intervals obtained from the two independent runs. For all these statistics, the effective sample size was greater than 100 and the relative discrepancy was less than 0.3.

## Ethics approval and consent to participate

Not applicable.

## Consent for publication

All authors agreed to submit this version of the manuscript.

## Availability of data and material

All datasets used in the research for the manuscript have been available via the following link: https://datadryad.org/stash/share/MqfshOJSWkeWWZZbgYsE7AqmaTYP9PpRtAjTXOtT1G8.

## Authors’ contributions

PB designed the protocol, supervised and contributed to the analysis of data. MM collected, analysed and interpreted data and prepared the first draft of the manuscript. PB and MM discussed the results, edited the final version of the manuscript and gave final approval for publication.

## Acknowledgments

The authors thanks Laure Devine for her comments and corrections on the manuscript, and Dany Lemay for his help with the computer aspects of the analysis.

